# Different states of priority recruit different neural codes in visual working memory

**DOI:** 10.1101/334920

**Authors:** Qing Yu, Bradley R. Postle

## Abstract

We tracked the neural representation of the orientation and the location of stimuli held in working memory at different levels of priority (“attended” and “unattended” memory items -- AMI and UMI), using multivariate inverted encoding models of human fMRI. Although representation of the orientation of the AMI and of the UMI could be reconstructed in several brain regions, including in early visual and parietal regions, the identity of the UMI was actively represented in early visual cortex in a distinct “reversed” code, suggesting this region as a site of the focus of attention to nonspatial stimulus information. The location of stimuli was also broadly represented, although only in parietal cortex was the location of the UMI represented in a reversed code. Our results suggest that a common recoding operation may be engaged, across stimulus dimensions and brain areas, to retain information in working memory while outside the focus of attention.

## Introduction

Important for understanding the flexible control of behavior [1, 2] is understanding working memory, the mental retention of task-relevant information and the ability to manipulate it and use this information to guide contextually appropriate actions [3, 4]. State-based theoretical models of working memory posit that information can be held at different levels of priority in working memory, with information at the highest level of priority in the focus of attention (FoA), and the remaining information in a variously named state of “activated long-term memory” [5] or “region of direct access” [6]. Much of the empirical support for these models comes from tasks using a “retrocuing” procedure in which, after a trial’s to-be-remembered information has been removed from view, a subset of that information is cued to indicate that it will be tested. Retrocuing can both improve memory performance behaviorally [7] and increase the strength of retrocued information neutrally [8].

The retrocuing procedure allows for the controlled study of the back-and-forth switching of priority between memory items that is required for many complicated working memory tasks, such as the n-back [9] and working memory span [10] tasks. In the dual serial retrocuing (DSR) task, two items are initially presented as memoranda, followed by a retrocue that designates one the “attended memory item” (AMI) that will be interrogated by the impending probe. The uncued item cannot be dropped from working memory, however, because following the initial memory probe, a second retrocue may indicate (with *p* = 0.5) that this initially uncued item will be tested by the second memory probe. Thus, following the initial retrocue, the uncued item becomes an “unattended memory item” (UMI) [11]. Functional magnetic resonance imaging (fMRI) and Electroencephalography (EEG) studies of the DSR task have demonstrated that an active representation was only observed for the AMI, but not for the UMI, using multivariate pattern classification (MVPA) [12–14]. Thus, an elevated level of activation, particularly in temporo-occipital networks associated with visual perception, may be a neural correlate of the FoA. The neural bases of the UMI, however, are less clear.

Most DSR studies to date have failed to find MVPA evidence for an active representation of the UMI [12–14], although such a trace can be transiently reactivated with a pulse of transcranial magnetic stimulation (TMS) [15]. The one study that has found evidence for active representations of the UMI localized them to intraparietal sulcus (IPS) and the frontal eye fields (FEF), in an analysis of fMRI data from 87 subjects [16]. Thus, the current preponderance of extant data suggests that the neural representation of the UMI may be at a level of sustained activity that is so low as to be at or below the boundary of what can be detected with current methods and conventional set sizes. Although there are mechanisms other than elevated activity that could represent information in working memory [17, 18], the work presented here was designed to assess two alternative hypotheses about the neural representation of the UMI that have received less attention to date. One is that the representation of the UMI may be active, but in a representational format fundamentally different from the AMI, and therefore difficult to detect with MVPA methods. The second is that what may be most prominently maintained in working memory is a representation of the trial-unique context in which the UMI was presented, rather than a representation of stimulus identity per se.

Although MVPA decoding approaches are powerful analytic techniques that can provide evidence of whether two kinds of information are different, they are inherently limited in that they don’t directly provide information about how they differ. Therefore, in the current study we used multivariate inverted encoding modeling (IEM) [19–22] to evaluate item-level mnemonic representations of AMIs and UMIs. By specifying an explicit model of how stimulus properties are represented in large populations of voxels, we could assess quantitative and qualitative changes in stimulus representation as a function of changes in priority status. IEM may also be a more sensitive method for tracking working memory representations [22].

Our results revealed two important properties of UMI representations: first, rather than being just a “weak AMI”, the orientation of the UMI is actively represented in early visual cortex in a format that is different from the AMI; second, contextual information is also supported by a similar recoding scheme; that is, parietal cortex maintains a representation of the location of both memory items that also encodes their priority status, a property absent from spatial representations in early visual cortex.

## Results

### Experiment 1

#### Behavioral results

Participants performed two DSR tasks (*Retain1* and *Retain2*) in the scanner. In the *Retain1* task, although two orientation patches were initially presented as targets, the same one was always cued twice, meaning that the initially cued orientation remained in the focus of attention (i.e., the AMI) for the remainder of the trial, and the uncued item could be dropped from memory (“dropped memory item,” DMI). In the *Retain2* task, the initially uncued item became a UMI, because it was possible that it would be cued by the second retrocue (Fig 1A). Accuracies of responses to two probes were averaged to assess the behavioral performance of each task. Accuracy in the *Retain1* (67.4% ± 2.4%) and *Retain2* (68.8% ± 1.4%) tasks did not differ significantly (*t*(9) = 0.758,*p* = 0.468), nor did accuracies for the Stay (69.3% ± 1.5%) and Switch (68.2% ± 1.6%) conditions of the *Retain2* task (*t*(9) = 0.749, *p* = 0.473).

**Fig 1.**
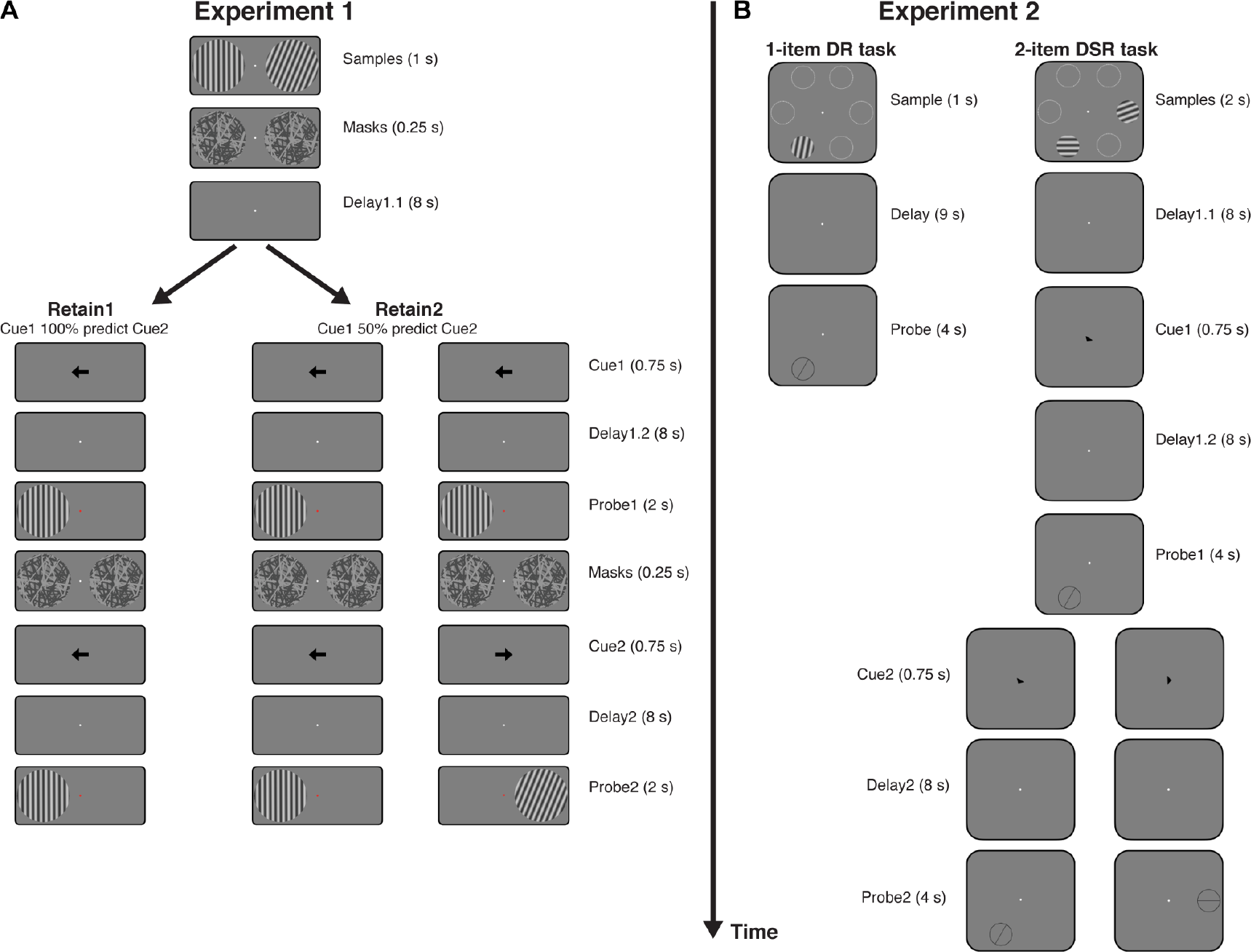
Experimental procedure. **A**. In Experiment 1, participants performed two tasks in the scanner. In *Retain1* task, participants remembered two orientations for Delay1.1, one in each hemifield, and were cued on one of them for Delay1.2. After a first probe, the same cue appeared and participants needed to recall the same orientation once again after Delay2. The probe task was a change detection task. In *Retain2* task, participants underwent the same procedure, except that the second cue may switch to the other orientation on 50% of the trials. **B**. In Experiment 2, participants performed the *Retain2* task only, and the two orientations could appear in two of six different locations (white circles are for demonstration purposes and were not present during the actual experiment). Participants performed a delay-estimation task of orientations. Besides the main experiment task, participants also performed a 1-item working memory task for training independent IEMs.

#### Reconstructing the neural representation of orientation of the AMI, the DMI, and the UMI

Our analytic strategy was to compare IEM reconstructions from models trained on different memory items to assess the similarity of representational format between the trained and tested items. Different IEMs were trained using the AMI, the UMI, or the DMI labels. Training was performed for the *Retain1* and *Retain2* tasks separately. For AMI-trained IEMs, the AMI labels were used to train the IEM, and the IEM was tested on data from both AMI-labeled and UMI/DMI-labeled data. When tested with UMI/DMI-labeled data, reconstructions from this AMI-trained IEM would index the extent to which the representational format of the UMI/DMI was similar to that of the AMI. For UMI/DMI-trained IEMs, the IEM was trained on the UMI/DMI and tested on both the AMI and the UMI/DMI. This IEM allowed us to examine the UMI/DMI representation without assuming any relationship with the representational format of information in the focus of attention.

For Experiment 1 we focused on an early visual ROI (V1-V2), because many studies have found robust evidence for an active representation of the AMI in this brain region. Furthermore, no studies, including the study finding evidence for an active representation of the UMI [16], have found evidence for an active representation of the UMI in this region. All the *p*-values were corrected across tasks and conditions using False Discovery Rate (FDR) method in this and subsequent analyses.

We focused on the TR 18 s after trial onset in order to maximize our ability to see the effect of the retrocue on stimulus representation during Delay1.2. In the *Retain1* task, whereas the orientation of the AMI could be reconstructed with the AMI-trained IEM (*p* = 0.031), reconstruction of the DMI was unsuccessful, whether tested with the AMI-trained or the DMI-trained IEM (*p*s = 0.354 and 0.665). In the *Retain2* task, reconstructions of the orientation of the AMI and of the UMI with the AMI-trained IEM went in opposite directions: a positive reconstruction approaching significance for the AMI (*p* = 0.084) and a significant negative reconstruction for the UMI (*p* = 0.011). The negative reconstruction of the UMI had the lowest response in the target channel, and progressively higher responses in non-target channels that grew with the distance of the non-target channel increased (Fig 2). The UMI could not be reconstructed with a UMI-trained IEM (*p* = 0.212), although a marginally significant negative reconstruction for the AMI was observed with this UMI-trained IEM (*p* = 0.085; S1 Fig).

**Fig 2.**
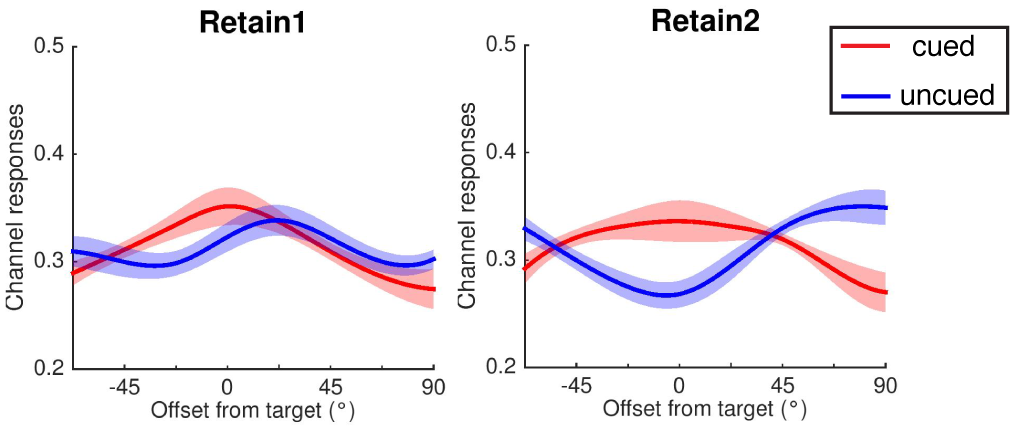
Experiment 1: Orientation reconstruction during Delay1.2 in the early visual ROI. Orientation reconstructions in the *Retain1* and *Retain2* tasks in Delay1.2. Red line represents the cued orientation (AMI during Delay1.2), and blue line represents the uncued orientation (DM1 for *Retain1* task and UMI for *Retain2* task during Delay1.2). Reconstructions were averaged across all participants. Continuous curves were created with spline interpolation method for demonstration purposes. Channel responses are estimated BOLD responses in relative amplitude. Shaded areas indicate ± 1 SEM.

The finding of a reliable negative reconstruction for the UMI during Delay1.2 was noteworthy because it deviated from the expectations that we would either replicate previous failures to find evidence for an active representation of the UMI during Delay1.2 [12–15], or we would find that the reconstruction of the UMI was qualitatively the same as that of the AMI, but simply lower in magnitude. Instead, the negative reconstruction that we observed suggested a distributed pattern of activity that differed both from the trained pattern of the AMI and from baseline, implying an active representation with a code that was different from the code with which the AMI was represented during Delay1.2. Furthermore, this finding would implicate early visual cortex in the active representation of the UMI, which would be at variance with accounts positing a privileged role for higher-level regions in visual working memory storage during conditions involving shifting attention [16] or distraction [23, 24]. Finally, this finding would represent, to our knowledge, the first report of a negative IEM reconstruction as an interpretable index of the state of an *active* neural representation of stimulus information.

### Experiment 2

Due to its novel and unexpected nature, it was important that we replicate the evidence from Experiment 1 for an *active* but *negative* representation of the UMI in early visual cortex. With Experiment 2, we also sought to extend this finding in several ways. First, we would extend our analyses into parietal regions that have also been implicated in the working memory representation of information, specifically, to the IPS. Second, we would investigate in greater detail the representational bases of the UMI by training IEMs with data from a variety of cognitive conditions. Finally, we would investigate how the representation of an item’s trial-specific context – i.e., its location on the screen -- might be differently sensitive to changing priority. To elaborate, to succeed on the DSR task requires not only a memory of the orientation of the two items presented at the beginning of the trial (say, a patch with an orientation of 20° and another with an orientation of 120°), but also a memory of *where* on the screen each of these items has been presented. We have hypothesized that, because maintaining the binding between an item’s identity and its context is necessary to keep it in working memory [25, 26], this contextual information may be represented in a parietal salience map [Gosseries, Yu, et al., 201827]. Therefore, we designed Experiment 2 to also assess the mnemonic representation of location context by modifying the DSR to feature 6 possible locations at which the two orientation patches could be presented on any trial.

#### Behavioral results

Experiment 2 required recall responses, which were fit with a 3-factor mixture model that estimated the precision of responses (“concentration”), the probabilities of responses to the target (*p*T), of responses to a nontarget (*p*N), and of guess responses (see Methods). Performance on the Stay vs. Switch conditions was marginally different in terms of concentration (16.93 ± 2.74 for Stay vs. 11.35 ± 1.67 for Switch; *t*(9) = 2.211, *p* = 0.054), and did not differ along the other parameters (*p*T: 79.9% ± 1.9% vs. 76.3% ± 3.1%; *p*N: 3.7% ± 1.7% vs. 4.9% ± 2.4%; and guessing: 16.4% ± 1.9% vs. 18.8% ± 2.9%), *t*s < 1.199, *p*s > 0.261.

#### Reconstructing representations of the orientation of the AMI and UMI

In addition to AMI- and UMI-trained IEMs, replicating the procedure from Experiment 1, for Experiment 2 we also trained IEMs on an independent 1-item delayed recall (DR) task, so as to more thoroughly evaluate the representational bases of memory items. IEMs trained on 1-item DR task would provide “idealized” estimates of how the brain represents stimulus orientation and location when only a single item is being processed, thereby allowing us to train IEMs unconfounded by factors that may be associated with processing two stimuli simultaneously. Additionally, 1-item DR-trained IEMs would be independent of the DSR task, and so would support direct comparisons of IEM reconstructions between different conditions in the DSR task. *P*-values reported in this section were corrected across conditions and ROIs.

##### AMI- and UMI-trained IEMs

We first sought to replicate the findings in Experiment 1 by repeating the analyses in Experiment 1 on Experiment 2. IEMs were trained and tested on the same late-Delay1.2 TR (18 s after trial onset) as in Experiment 1. In early visual cortex (V1-V2), patterns of reconstructions of orientation were broadly similar to the findings from Experiment 1 (Fig 3A): AMI-trained IEM produced significantly positive reconstruction of the AMI in late Delay1.2 (*p* = 0.011), and significantly negative reconstruction of the UMI (*p* = 0.046); furthermore, UMI-trained IEM failed to reconstruct the UMI (*p* = 0.330). In IPS, however, we observed a qualitatively different pattern (Fig 3A): a robust positive reconstruction of the orientation of the AMI (*p* = 0.042) was accompanied by a marginally positive reconstruction of the orientation of the UMI (*p* = 0.063). Also at variance with the early visual ROI, with a UMI-trained IEM in IPS we could successfully reconstruct the UMI (*p* = 0.014) as well as the AMI at a statistically marginal level (*p* = 0.088; S2A Fig). No significant difference was observed between the two reconstructions in IPS for either IEM (both *p*s > 0.505). Together, these results indicated that although the UMI could be reconstructed in both early visual cortex (replicating Experiment 1) and in IPS, it was represented in different formats in these two regions – different from the AMI in early visual cortex, similar to the AMI in IPS.

**Fig 3.**
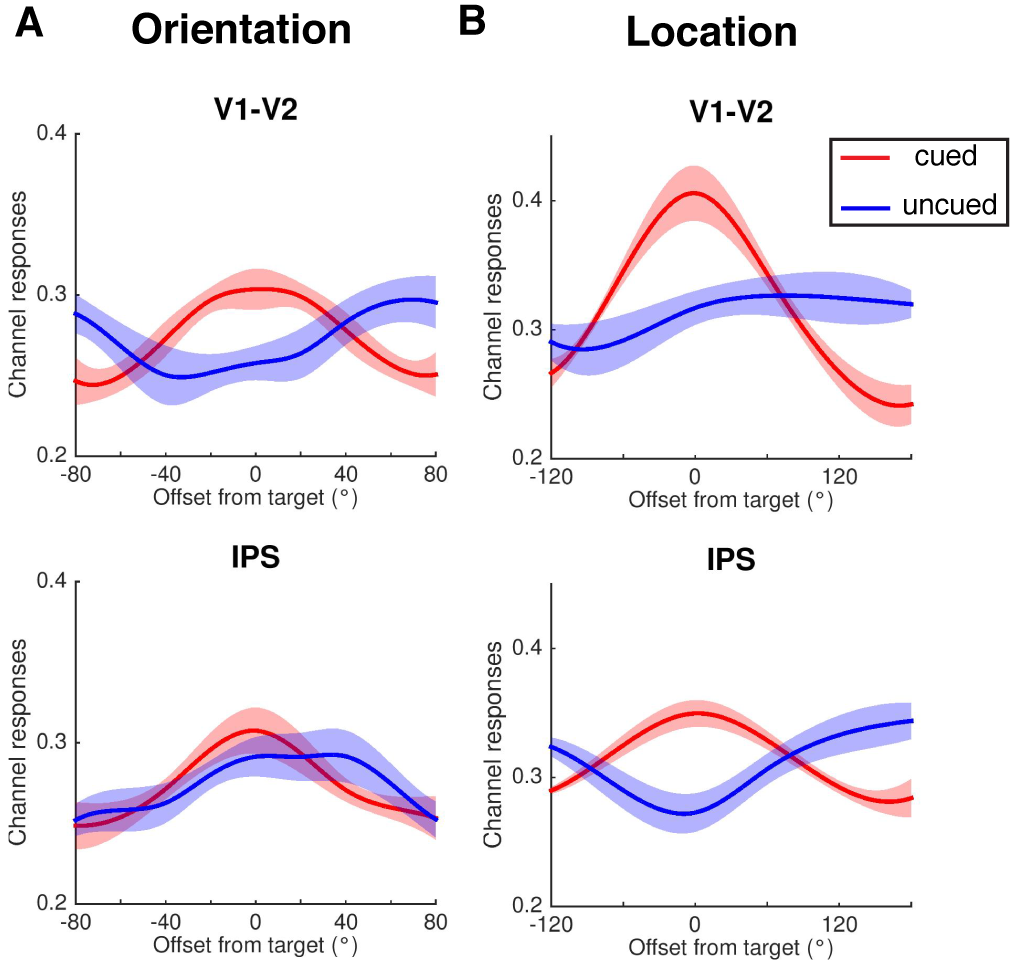
Experiment 2: Orientation and location reconstructions in early visual ROI and IPS. Orientation (**A**) and location (**B**) reconstructions during Delay1.2 in two representative ROIs: V1-V2 (early visual cortex) and IPS (parietal cortex), using an AMI-trained model. Red line represents the cued orientation (AMI during Delay1.2), and blue line represents the uncued orientation (UMI during Delay1.2). Reconstructions were averaged across all participants. Continuous curves were created with spline interpolation method for demonstration purposes. Channel responses are estimated BOLD responses in relative amplitude. Shaded areas indicate ± 1 SEM.

##### 1-item DR-trained IEM

Next, we sought to reconstruct the orientations of the AMI and UMI using a model trained with data from the TR beginning 4 s after sample onset of the 1-item DR task (i.e., sample-evoked signal), and tested on the TR from late Delay1.2 (18 s after sample onset) of the DSR task. In the early visual ROI, reconstruction of the AMI was robust (*p* = 0.008). For the UMI in the same ROI, reconstruction of the UMI was also positive (*p* < 0.00001). Similarly, in IPS, reconstructions of the AMI and of the UMI with the 1-item DR-trained IEM were both significantly positive (*p*s = 0.028 and 0.002; S3A Fig). Moreover, the two reconstructions were not significantly different from each other, in either early visual cortex or IPS (both *p*s > 0.422).

#### Reconstructing representations of the location of the AMI and UMI

##### AMI- and UMI-trained IEMs

To reconstruct the location where the AMI and the UMI had been presented, we first used the leave-one-run-out procedure with signal from the late Delay1.2 TR (18 s after trial onset). In early visual cortex, whereas the location of the AMI could be reconstructed with an AMI-trained IEM (*p* < 0.00001), the location of UMI could not (*p* = 0.910; Fig 3B). In IPS, in contrast, location of both the AMI and of the UMI could be reconstructed, with a positive reconstruction of the location of the AMI (*p* = 0.0008) and a negative reconstruction of the location of the UMI (*p* = 0.023; Fig 3B).

With UMI-trained IEMs, in stark contrast to what was observed for reconstruction of orientation, the location of the UMI could be reconstructed both in early visual cortex and in IPS (*p*s = 0.007 and 0.038; S2B Fig).

##### 1-item DR-trained IEM

For reconstructions with IEMs from the independent 1-item DR task we trained two IEMs -- one with data from the TR beginning 4 s after sample onset (i.e., sample-evoked signal), and one with data from the TR beginning 10 s after sample onset (i.e., delay-period signal). In the early visual ROI, the location of both AMI and UMI could be successfully reconstructed with both IEMs (*p* < 0.00001 and *p* = 0.002, respectively, for the sample-trained IEM; and *p* = 0.001 and *p* < 0.00001, respectively, for the delay-trained IEM). In IPS, in contrast, reconstruction attempts were unsuccessful for all items with both IEMs (*p*s > 0.139; S3B Fig and S4 Fig).

#### Validation of the negative reconstructions

To validate the negative reconstructions observed in Experiments 1 and 2, we took several steps to explore possible artifactual explanations for these results. First, we considered the possibility that the negative reconstruction of the orientation of the UMI may have reflected influences from the AMI, because the two could never take the same value on the same trial, but instead always had a minimal distance of separation: at least 22.5° of orientation in Experiment 1; at least 20° of orientation in Experiment 2; and at least 60° of polar angle for location in Experiment 2. The reasoning behind this alternative account is that recentering all UMI reconstructions on a common target channel would necessarily produce a situation in which every AMI fell on a non-target channel, and this could result in a negative-going reconstruction after averaging across trials. One reason to doubt this alternative account a priori is that the average angular distance between the AMI and the UMI across all trials was relatively small. For example, for orientations in Experiment 2, the average angular distance was 49.5° ± 2.0° across participants. In contrast, if the minimum distance constraint between the two orientations did not exist, and the two sample orientations on each trial were randomly sampled from 0° to 180°, the average angular distance would be ~45°. It seems unlikely that this ~5° of difference could be responsible for what would need to be a roughly 45° bias in UMI reconstructions, the size of bias that would be needed to produce artifactual negative-looking reconstructions of the kind that we observed. Furthermore, a negative reconstruction was not observed for the DMI in the *Retain1* task in Experiment 1, despite the fact that its procedural conditions were identical.

A second way to assess the minimal-angle-of-separation alternative account analytically is to compare UMI representations with representations of “absent” orientations/locations. To do this, on each trial, we randomly selected an orientation/location that had not been presented, reasoning that this absent orientation/location also had a minimum angular distance from the AMI, and attempted its reconstruction with the same procedures that generated negative reconstructions of UMI. This exercise generated two important results: First, none of the reconstructions of an item absent from the trial’s memory set was significant (all *p*s > 0.893); and second, the negative reconstructions of UMI orientation in Experiment 1 and of UMI of location in Experiment 2 were significantly different (i.e., “more negative”) from reconstructions of absent items (*p*s = 0.012 and 0.028, respectively).

As a third control, we randomly permuted the UMI labels across trials within each participant, a procedure that ensured that the mean orientation difference between the AMI and the UMI across trials remained constant. Reconstructions with permuted UMI labels were not significant either (all *p*s > 0.981). Additionally, negative reconstructions of UMI orientation in Experiment 1 and of UMI location in Experiment 2 were also significantly “more negative” than reconstructions of permuted UMIs (*p*s = 0.018 and 0.031, respectively).

As a fourth control, we performed the AMI-trained IEM analyses on a simulated dataset in which the response in each voxel was determined by its preferred orientation and random Gaussian noise. The rationale was that if the simulated data only contained information about the AMI, then recentering the reconstructions by arbitrarily generated UMI labels would not result in either positive or negative reconstruction of the UMI. As expected, this simulated IEM resulted in robust reconstruction of the AMI (*p* < 0.00001), and no reconstruction of the UMI (*p* = 0.740), although the AMI and the UMI maintained a minimum orientation distance as in the actual experiments (S5 Fig). This result suggested that the negative UMI reconstruction observed in our experiments could not be explained by influence from the AMI. This simulation evidence, together with all the evidence above, suggest that the negative reconstruction of the UMI is reliable and cannot be simply explained by minimum distance between the AMI and the UMI.

Lastly, as a fifth control, we fitted the reconstructions of our primary results using an exponentiated cosine function to assess the significance of each reconstruction (instead of the slope of the reconstruction). The model fits yielded qualitatively similar results to those of slope measures (S1 Table), indicating that our results are consistent across different quantification methods.

## Discussion

Neural representations of information, including of information held in working memory, are thought to be supported by anatomically distributed networks [16, 22, 28, 29]. What remains unclear is the extent to which stimulus-related patterns of activity localized to different brain regions may employ the same or different representational formats, and may support similar or different functions. In the current study we manipulated the momentary state of priority of information in working memory, and employed multivariate encoding models to track interregional differences in the representation of behaviorally relevant information.

### Representation of stimulus identity in visual working memory

With regard to the representation of stimulus identity (here, orientation), our results indicate that early visual cortex supports the multi-dimensional representation of stimulus identity: although the representations of orientation of the AMI and of the UMI shared some features in common (in that both yielded similar reconstructions by an IEM trained on perception/encoding-related signal from a 1-item task); they also differed markedly (in that the reconstruction of the UMI was significantly different from that of the AMI when tested with the AMI-trained IEM). The results from IPS, in contrast, provided evidence for only a common representational format, in that none of the successful reconstructions of the AMI and of the UMI differed from each other.

The fact that information with different attentional priority is represented in a different representational format in early visual cortex, but not in IPS, supports the view that the former is an important site for the focus of attention in visual working memory, an observation consistent with sensorimotor-recruitment models of visual working memory [4, 30]. It also suggests that the brain may implement (at least) two distinct, but mutually compatible, schemes for the retention of UMIs: recoding stimulus information into multiple formats in early visual cortex, and maintaining stimulus representations in multiple anatomically distinct networks (as also suggested by previous studies: e.g., [16, 23, 28].

### Representation of location context in visual working memory

Although our DSR task explicitly tested visual working memory for a nonspatial stimulus feature, the task can nevertheless not be performed successfully without the trial-specific representation of the location at which each stimulus was presented. Indeed, context binding may be essential of working memory [25, 26]. Furthermore, many studies have demonstrated the automatic binding of location information to the to-be-remembered visual features [31–34].

Because delay-period BOLD signal intensity in IPS is markedly higher on trials that require visual working memory for 3 items drawn from the same category than for 3 items drawn from different categories [Gosseries, Yu, et al., 201827], it may be that IPS recruitment scales, at least in part, with demands on context binding. This would be consistent with the idea that a parietal salience map tracks the location context of all items held in visual working memory. In Experiment 2, the location representations of the AMI and of the UMI were differently sensitive to attentional priority in different brain regions. Although both early visual and IPS ROIs maintained representations of the locations of the AMI and of the UMI, early visual cortex did not support AMI-encoded representations of the location of the UMI; IPS, in contrast, simultaneously supported reverse-AMI-encoded representations of the location of the UMI, in addition to AMI-encoded representations of the AMI and UMI-encoded representations of the UMI. Thus, unlike early visual cortex, IPS represented the location of all items in working memory in a manner that encoded the priority status associated with each of those locations. Notably, this pattern of results is the reverse of what was observed for the differential representation of orientation in these two regions. The results from IPS are consistent with the possibility that context and priority in visual working memory are represented by the same parietal salience map that tracks these factors during behaviors that do not make any overt demands on working memory [35–37].

Furthermore, it is noteworthy that although representations of the location of the AMI and of the UMI in IPS were robust with regard to AMI- and UMI-trained models, they could not be reconstructed with the 1-item DR-trained model. This suggests the representation of location in IPS is highly dependent on task context (e.g., tracking the location of just one item vs. the location of two items simultaneously). Methodologically, it highlights the utility of incorporating results from different models when interpreting null results in multivariate analyses of stimulus representation in working memory.

### Negative reconstructions of the representation of orientation and of location context

The delay-period representation of stimulus information (a.k.a., “storage”) is a cardinal property of working memory. Although our results make clear that many brain areas can simultaneously represent information about the same stimulus dimension, often in similar representational formats, it seems unlikely that any two regions’ functions are completely redundant. Some previous work has also suggested that representations in different brain areas are differentially sensitive to attention, distraction, or other types of task manipulations [16, 24, 38, 39], as evidenced by a decline in decoding or encoding performance in some brain regions but not others. Here, we interpret our results as reflecting multiple graded distributions of functional activity, with the likelihood that, for some circuits and in some instances, the primary function being supported is one other than storage, per se. We propose that the recoding of stimulus orientation information into a reverse-AMI-encoded representation in occipital cortex, and of stimulus location information into a reverse-AMI-encoded representation in IPS, may reflect a mechanism for accomplishing the robust retention of information about stimuli that are in working memory but outside the focus of attention.

Why should the features of a stimulus be represented differently when it is a UMI than when it is an AMI? One possible explanation, deriving from information theory, follows from the observation that the requirement of temporarily storing information in a noisy neuronal network, for later retrieval, is mathematically equivalent to transmitting that information through a noisy channel [40]. Shannon [41] demonstrated that high-fidelity transmission of information though a noisy channel can be accomplished by recoding the message into a format that takes into account the structure of the noise, then decoding it at the receiving end. One possible account of the “negative reconstructions” that we have reported here, then, is that they may reflect a common computation that exploits this principle to maintain high-fidelity representations of stimulus information while it is held in working memory, but outside the FoA.

A second possible account of the reverse-encoding of the UMI is that it may reflect the projection of this information into a dynamical “null space” that keeps it active, but prevents it from being read-out by downstream systems that influence behavior. Such dynamics have been most closely studied in the motor system, in which high-dimensional motor preparatory activity is construed as a dynamic interplay between subpopulations whose aggregate output projects into a (high-dimensional) output-null space that simultaneously allows for active planning/preparation while preventing these same signals from influencing downstream effector systems [42]. A similar scheme has been proposed for the dynamics of delay-period activity in the PFC of the nonhuman primate [43]. Thus, in the results reported here, it may be that the reverse-encoded representations of the UMI serve to keep it in an “output-null” state.

We note that these instances of negative reconstruction can’t be characterized as inhibition, because the effect of inhibition would be to “flatten” an IEM reconstruction. Nor are they likely to be the inhibitory engrams postulated by Barron and colleagues [44], because whereas the effect of the inhibitory engram would be to minimize representation-related activity, the negative reconstructions that we have described here must be the result of an active reconfiguration of activity in all the voxels feeding into that IEM. Thus, although these reverse-AMI-encoded representations are, indeed, quantitatively “negative reconstructions”, in functional terms it may be more fitting to characterize them as “different from” the code on which the IEM was trained.

## Materials and Methods

### Participants

A total of twenty-two individuals participated in the study. Twelve individuals (5 males, mean age 23.3 ± 3.6 years) participated in Experiment 1. Two (both females) were excluded from analysis due to low behavioral performance on the memory task (average accuracy < 60%). Another ten individuals (4 males, mean age 23.8 ± 3.5 years) participated in Experiment 2. All were recruited from the University of Wisconsin-Madison community. All had normal or corrected-to-normal vision, were neurologically healthy, and provided written informed consent approved by the University of Wisconsin-Madison Health Sciences Institutional Review Board. All participants were monetarily compensated for their participation.

### Stimuli and Procedure

All stimuli were created and presented using Matlab (MathWorks, Natick, MA) and Psychtoolbox 3 extensions [45, 46] on a 60-Hz Avotec Silent Vision 6011 projector (Avotec, Stuart, FL), and viewed through a coil-mounted mirror in the MRI scanner. An fMRI-compatible button box (Experiment 1) or trackball response pad (Current Designs Inc., Philadelphia, PA; Experiment 2) was employed to record the behavioral responses.

#### Experiment 1

Participants performed two dual serial retrocuing (DSR) tasks (*Retain1* and *Retain2*) in the scanner. A white fixation dot was presented at the center of the screen throughout the experiment. During the *Retain1* trials, participants first viewed two sample stimuli (sinusoidal gratings: radius = 5°; contrast = 0.6; spatial frequency = 0.5 cycles/°; phase angle randomized between 0° and 180°), each with a different orientation, and presented simultaneously (one in each hemifield, and centered on the horizontal meridian at 7° from fixation) for 1 s. After an interval of 0.5 s, two masks composed of randomly oriented black and white lines were presented at the sample locations for 0.25 s, followed by the first delay period. After 8 s (“Delay1.1”), an arrow displayed at fixation for 0.75 s pointed to the one of the two locations that had been occupied by the samples to indicate which would be tested by the next memory probe (“Cue1”). After an additional 8 s (“Delay1.2”), a probe grating requiring a Y/N recognition response was presented for 0.5 s in the same location as the sample it was testing, followed by a response period of 1.5 s (“Probe1”). Probe 1 was followed by two masks that were identical to the first two masks (0.25 s in duration), a 0.5 s blank interval, an arrow identical to Cue1 (“Cue2;” 0.75 s), a delay of 8 s (“Delay2”), and a probe grating (“Probe 2”) identical in timing to Probe1. Both probes required a button press indicating whether orientation of the probe grating did or did not match the cued grating. Half of the probes had an orientation that matched that of the cued grating, and the other half had an orientation that differed by 10° to 20°. The color of the fixation dot was changed to red during probe until participants made a button press. Intertrial-interval was either 4 s or 6 s. *Retain2* trials had exactly the same procedure as *Retain1* trials, except that Cue1 did not predict Cue2. Therefore, on half of the trials, Cue2 was identical to Cue1, meaning that the same cued orientation would be probed twice (a *Stay* trial); and on the other half Cue2 was different from Cue1, meaning that Probe2 would probe memory for the target that had not been tested by Probe1 (a *Switch* trial, Fig 1A). Following our previous work, the item cued by Cue1 was termed the “attended memory item” (AMI); on *Retain1* trials the item that was not cued by Cue1 was called the “dropped memory item” (DMI, because it could be dropped from working memory), and on *Retain2* trials the item that was not cued by Cue1was termed the “unattended memory item” (UMI). The two tasks were administered in separate blocks and the order of the tasks was randomly chosen. Participants were informed which task they would be performing at the beginning of each run. On each trial, the orientations of the two samples were randomly selected from a fixed set of eight orientations (0°, 22.5°, 45°, 67.5°, 90°, 112.5°, 135°, 157.5°, with a random jitter between 0° and 3°), with the constraints that the same orientation could only be selected once for any given trial, and that each would be presented at each location once during each run. This resulted in a minimum distance of 22.5° between the two sample orientations on every trial. Each run began with an 8-s blank period, was comprised of 16 trials, and lasted 600 s. Eight of the participants performed six runs of the *Retain1* task, one performed seven runs and one performed twelve runs. All participants performed twelve runs of the *Retain2* task.

#### Experiment 2

Participants performed two working memory tasks in the scanner. A white fixation dot was presented at the center of the screen throughout the experiment. The first task was 1-item delayed recall (DR) task of orientation, intended for training IEMs that would be used to analyze data from this experiment’s DSR task. On each trial, one grating (radius = 2°, contrast = 0.6, spatial frequency = 0.5 cycles/°, phase angle randomized between 0° and 180°) was presented on the screen with an eccentricity of 7° and participants were asked to remember its orientation. The location of the grating was chosen from six fixed locations (60° of distance from each other, with two on the horizontal meridian), and the orientation of the grating was chosen from nine orientations (0°, 20°, 40°, 60°, 80°, 100°, 120°, 140°, 160°) with a random jitter between 0° and 3°. The grating appeared on the screen for 1 s, followed by a delay period of 9 s, and then by a response period of 4 s. During the response period, an orientation wheel (2° in radius) was presented at the same location as the sample grating, and participants needed to rotate the needle at the center of the wheel to make it match the remembered orientation as precisely as possible. The inter-trial-interval was fixed at 8 s. Each run consisted of eighteen trials, resulting in a run length of 404 s. Participants performed a total of 24 to 30 runs of the 1-item working memory task in two separate scan sessions.

The second task was a two-item DSR task that was similar to the *Retain2* task in Experiment 1, except that the two samples could appear in any of six possible locations (same locations as in the 1-item DR task). On each trial, participants viewed two gratings (parameters identical to those in the first task) presented at two of six fixed locations and were asked to remember both. The minimum distance between the two samples was 20° for orientation and 60° for location on every trial. The two gratings appeared on the screen for 2 s, followed by a first delay period (Delay1.1) of 8 s. After that a cue appeared at the center of the screen for 0.75 s, which was a triangle-shaped arrow that pointed to one of the two sample locations. After another 8 s (Delay1.2), an orientation wheel was presented at the same location as the cued grating, and participants needed to reproduce the cued orientation on the wheel within a 4-s response window. 0.5 s after the first response period, participants saw a second cue, 50% of which would point to the first cued location (Stay), and the other 50% would point to the first uncued location (Switch). After a third 8 s of delay (Delay2), a second orientation wheel was presented at the same location as the second-cued grating, and again participants needed to reproduce the cued orientation on the wheel in 4 s (Fig 1B). The inter-trial-interval was fixed at 8 s. Each run consisted of twelve trials, resulting in a run length of 536 s. Participants performed 12 runs of this DSR task in one scan session.

In both experiments, electrooculography (EOG) of vertical and horizontal eye movements was recorded using the BrainVision EEG system (BrainVision LLC, Morrisville, NC), while participants performed the tasks in the scanner to ensure central fixation throughout each trial.

### Behavioral analysis for Experiment 2

We analyzed behavioral responses with a three-factor mixture model [47] that uses maximum likelihood estimation to generate estimates of 1) the proportion of responses based on a representation of the probed item (“responses to target”); 2) the proportion of responses incorrectly based on a representation of the unprobed item (i.e., “misbinding” or “swap” errors); and 3) the proportion of responses that were guesses not based on either memory item; as well as 4) a “concentration” parameter that estimates the precision of target responses. Conceptually, the concentration parameter is similar to a model-free measure of the precision of responses that is computed as the inverse of the standard deviation of the distribution of responses.

### fMRI Data acquisition

Whole-brain images were acquired using a 3 Tesla GE MR scanner (Discovery MR750; GE Healthcare, Chicago, IL) at the Lane Neuroimaging Laboratory at the University of Wisconsin-Madison HealthEmotions Research Institute (Department of Psychiatry). Functional imaging was conducted using a gradient-echo echo-planar sequence (2 s repetition time (TR), 22 ms echo time (TE), 60° flip angle) within a 64 × 64 matrix (42 axial slices, 3 mm isotropic). A high-resolution T1 image was also acquired for each session with a fast spoiled gradient-recalled-echo sequence (8.2 ms TR, 3.2 ms TE, 12° flip angle, 176 axial slices, 256 × 256 in-plane, 1.0 mm isotropic).

### fMRI Data preprocessing

Functional MRI data were preprocessed using AFNI (http://afni.nimh.nih.gov) [48]. The data were first registered to the final volume of each scan, and then to anatomical images of the first scan session. Six nuisance regressors were included in GLMs to account for head motion artifacts in six different directions. The data were then motion corrected, detrended, and z-score normalized within each run.

### fMRI ROI definition

Anatomical ROIs were created by extracting masks from the probabilistic atlas of Wang and colleagues [49], and warping them to each subject’s structural scan in native space. Analyses in Experiment 1 were carried out in an early visual ROI (V1-V2 merged); analyses in Experiment 2 were carried out in an early visual ROI (V1-V2 merged) and a parietal ROI (IPS0-5 merged). All the ROIs were collapsed over the right and left hemispheres.

### Multivariate inverted encoding modeling

All inverted encoding model (IEM) analyses were performed using custom functions in Matlab. We used inverted encoding models (IEMs) to evaluate the representation of orientation (in Experiments 1 and 2) and of location (in Experiment 2) of the AMI and UMI. The IEM assumes that the responses of each voxel can be characterized by a small number of hypothesized tuning channels. In Experiment 1 the number of orientation tuning channels was eight, and in Experiment 2 the number of orientation tuning channels was nine and the number of location tuning channels was six. Following previous work [22, 29], the idealized feature tuning curve of each channel was defined as a half-wave-rectified and squared sinusoid raised to the sixth power (FWHM = 0.94 rad) for orientation in Experiment 1, to the eighth power (FWHM = 0.82 rad) for orientation in Experiment 2, and to the sixth power (FWHM = 1.88 in rad) for location in Experiment 2.

Before feeding the preprocessed data into the IEM, a baseline from each voxel’s response was removed in each run using the following equation from [19, 39]:

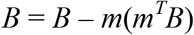

in which *B* represented the data matrix from each run with size *ν* × *c* (*ν*: the number of voxels in the ROI; *c*: the number of orientations/locations) and *m* represented the mean response across all stimulus conditions of length *ν*. A constant of 100 was added to *B* to avoid matrix inversion problems after baseline removal.

We then computed the weight matrix (*W*) that projects the hypothesized channel responses (*C*_*1*_) to actual measured fMRI signals in the training dataset (*B*_*1*_), and extracted the estimated channel responses (*Ĉ*_*2*_) for the test dataset (*B*_*2*_) using this weight matrix.

The relationship between the training dataset (*B*_*1*_, *ν* × *n*, *n*: the number of repeated measurements) and the channel responses (*C*_*1*_, *k* × *n*) was characterized by:

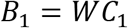

Where *W* was the weight matrix (*ν* × *k*).

Therefore, the least-squared estimate of the weight matrix 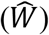 was calculated using linear regression:

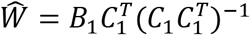

The channel responses (*Ĉ*_*2*_) for the test dataset (*B*_*2*_) was then estimated using the weight matrix 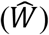:

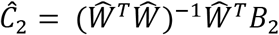

For Experiment 1, we used a leave-one-run-out procedure to build the weight matrix and to calculate the estimated channel outputs for orientations in the test dataset. IEMs were trained with signals from the time point of interest, for each task separately. Specifically, we used the TR 18 s after trial onset (7.5 s after Cue1 offset) to reconstruct the representation of orientation in Delay1.2. For *Retain1* task, the training orientation labels could either be the AMI (AMI-trained IEM) or the DMI (DMI-trained IEM); for *Retain2* task, the training orientation labels could either be the AMI (AMI-trained IEM) or the UMI (UMI-trained IEM). The obtained weight matrices were applied to the same time points in the test dataset. The estimated channel outputs obtained after each iteration were shifted to a common center, with 0° corresponding to the test orientation channel. Again, for *Retain1* task the test orientations could either be the AMI or the DMI; and for *Retain2* task the test orientations could either be the AMI or the UMI. The shifted channel outputs were then averaged across all iterations within each task in each participant. As a result, for each task and each IEM, we could obtain two orientation reconstructions, one for the AMI, and one for the DMI or UMI. For Experiment 2, the AMI-trained and UMI-trained IEMs were constructed using the same method as in Experiment 1. We again used the TR 18 s after trial onset (7.25 s after Cue1 offset) for the leave-one-run-out procedure. Additionally, we trained “independent” IEMs with data from the 1-item DR task, and tested these IEMs on data from the DSR task. We used the TR 4 s after trial onset to train an orientation IEM, and the TR 4 s or 10 s after trial onset to train a location IEM. All the IEMs were estimated for orientations and locations separately.

### Statistical analyses

To characterize the strength of each reconstruction, we collapsed over the channel responses on both sides of the target channel, averaged them, and calculated the slope of each collapsed reconstruction using linear regression [31, 50]. A larger positive slope indicates stronger positive representation, and a larger negative slope indicates stronger negative representation. We used a bootstrapping procedure [22] to characterize the significance of the slopes. For each task/ROI, ten orientation/location reconstructions were randomly sampled with replacement from the reconstruction pool of ten participants and averaged. This procedure was repeated 10000 times, resulting in 10000 average orientation/location reconstructions for each task/ROI, and correspondingly 10000 slopes. To obtain a two-tailed measure of the *p*-values, the probabilities of obtaining a positive (*p*_pos_) or negative (*p*_neg_) slope among the 10000 slopes was calculated separately, and the *p*-value of the bootstrapping test was calculated using the following equation:

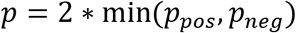

To characterize the difference between two slopes, we first calculated the difference between two bootstrapped slopes for 10000 times, which generated 10000 slope differences. The significance of the slope difference was then calculated using the same two-tailed method as above.

To compare the UMI reconstruction against reconstructions of “absent” orientations/locations, we generated a null distribution of “absent” orientation/location reconstructions by aligning the reconstruction centers to randomly selected absent orientations/locations (i.e., one orientation/location from the orientation/location pool that was neither the AMI nor the UMI). This procedure was repeated 10000 times for each task/ROI, and the obtained reconstructions were averaged across participants on each iteration. The robustness of absent feature reconstructions was evaluated using the same p-value calculation method (two-tailed) as mentioned in the previous paragraph. Moreover, we performed a more rigorous evaluation, in which the slope of the UMI was compared with the slope distribution of absent orientations/locations, and the probability of obtaining a larger UMI slope was computed (one-tailed). A significant *p*-value would indicate significantly negative reconstructions of the UMI compared to absent orientations/locations.

A similar procedure was conducted to compare the UMI reconstruction against reconstructions of permuted UMIs. The UMI labels were randomly shuffled across trials, and a null distribution of permuted UMIs was generated using the aforementioned method. Similarly, the robustness of permuted UMI reconstructions as well as the difference between UMI and permuted UMI reconstruction were evaluated.

To assess the consistency of our results across different quantification methods, we fitted an exponentiated cosine function to each reconstruction instead of slope:

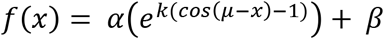

Where *x* is the channel responses, *α* corresponds to the amplitude, *β* corresponds to the baseline, *k* corresponds to the concentration, and *μ* corresponds to the offset (fixed at 0). The model fitting was performed by combining a general linear model with a grid search procedure [22]. First, we created a grid of parameters by defining 300 *k* values (between 0.1 and 30 in 0.1 steps). For each *k* we used the exponentiated cosine function (*α* = 1, *β* = 0, *μ* = 0) to generate a response function. The response function was then combined with a vector of ones (constant term) for estimation of *α* (regression coefficients) and *β* (constant term) using ordinary least squared regression method. We chose the set of parameters that had the minimum sum of squared errors as the best fits. We used a to quantify the robustness of reconstructions; positive and negative αs indicate positive and negative reconstructions, respectively. A two-tailed *p*-value was calculated for each reconstruction.

### Simulation of IEM reconstructions

We simulated a dataset of 700 voxels. Each voxel was assigned a preferred orientation, and the 700 preferred orientations were evenly spaced between 0° and 180°. Voxel response to the preferred orientation was a Gaussian function (amplitude = 1, baseline = 0, standard deviation = 0.25) and was characterized by responses in eight hypothesized orientation channels (0°, 22.5°, 45°, 67.5°, 90°, 112.5°, 135°, 157.5°). Random Gaussian noise with a standard deviation of 6 was added to the response in each voxel. We simulated 16 trials for each of the eight orientations, and these orientations served as the AMI labels. We then generated a same number of arbitrary UMI labels, following the same rules in our experimental task. As a result, the simulated AMI and UMI maintained the same minimum distance as in the actual experiments. The IEM was trained with simulated AMI labels, and this simulated AMI-trained IEM was then tested on both the simulated labels of the AMI and of the UMI. This simulation was repeated ten times and each simulated result was treated as the result of a single participant. The robustness of simulated reconstructions across ten simulations was characterized using the same bootstrapping method as in the actual experiments.

## Acknowledgements

This work was supported by National Institutes of Health grant R01MH064498 to B.R.P.

## Competing Interests statement

The authors declare no competing interests.

## Supporting Information

**S1 Fig.**
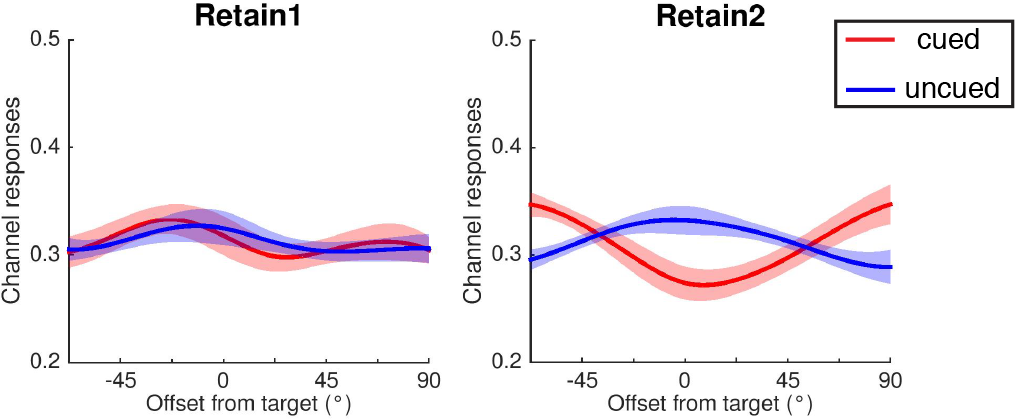
Experiment 1: Orientation reconstructions from a DMI-trained IEM (for *Retain1* task) and a UMI-trained IEM (for *Retain2* task). Red line represents the cued orientation (AMI during Delay1.2), and blue line represents the uncued orientation (DMI for *Retain1* task and UMI for *Retain2* task during Delay1.2). Reconstructions were averaged across all participants. Continuous curves were created with spline interpolation method for demonstration purposes. Channel responses are estimated BOLD responses in relative amplitude. Shaded areas indicate ± 1 SEM.

**S2 Fig.**
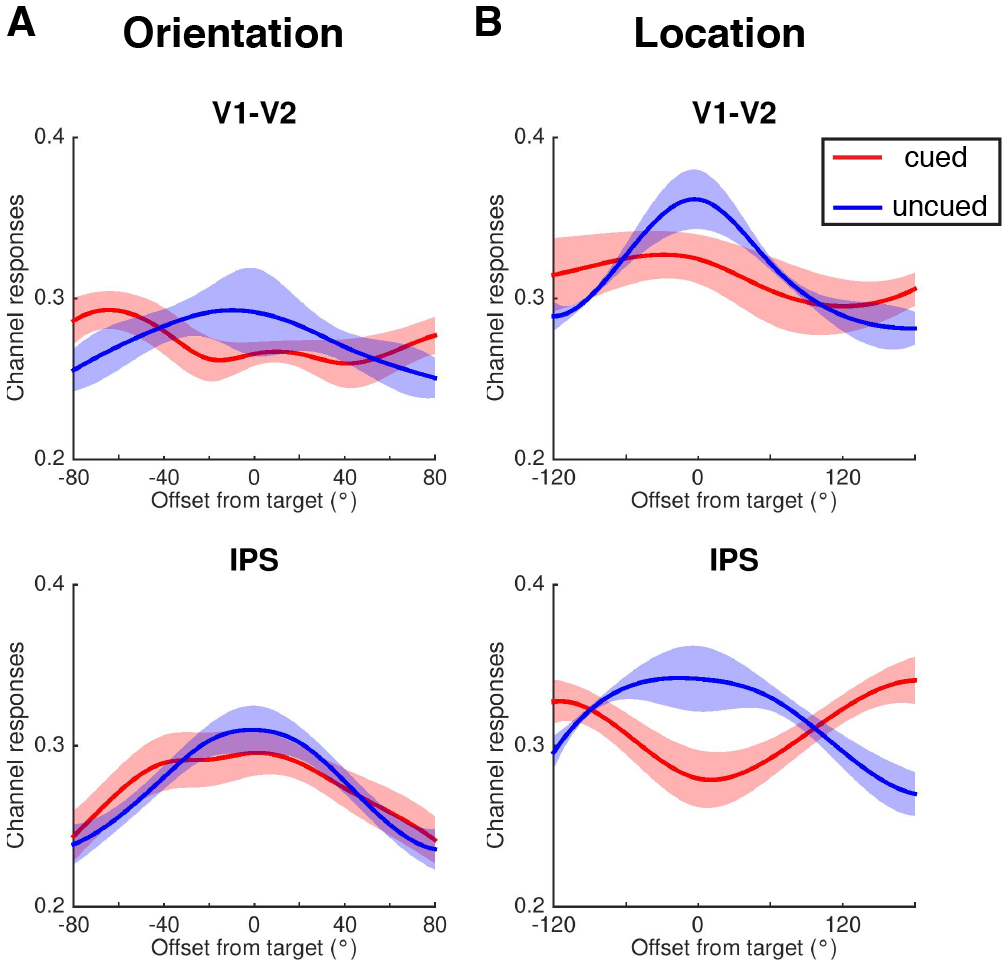
Experiment 2: Orientation and location reconstructions from a UMI-trained IEM, in early visual ROI and in IPS. Orientation (**A**) and location (**B**) reconstructions during Delay1.2 in V1-V2 (early visual cortex) and IPS (parietal cortex), using an UMI-trained model. Red line represents the cued orientation (AMI during Delay1.2), and blue line represents the uncued orientation (UMI during Delay1.2). Reconstructions were averaged across all participants. Continuous curves were created with spline interpolation method for demonstration purposes. Channel responses are estimated BOLD responses in relative amplitude. Shaded areas indicate ± 1 SEM.

**S3 Fig.**
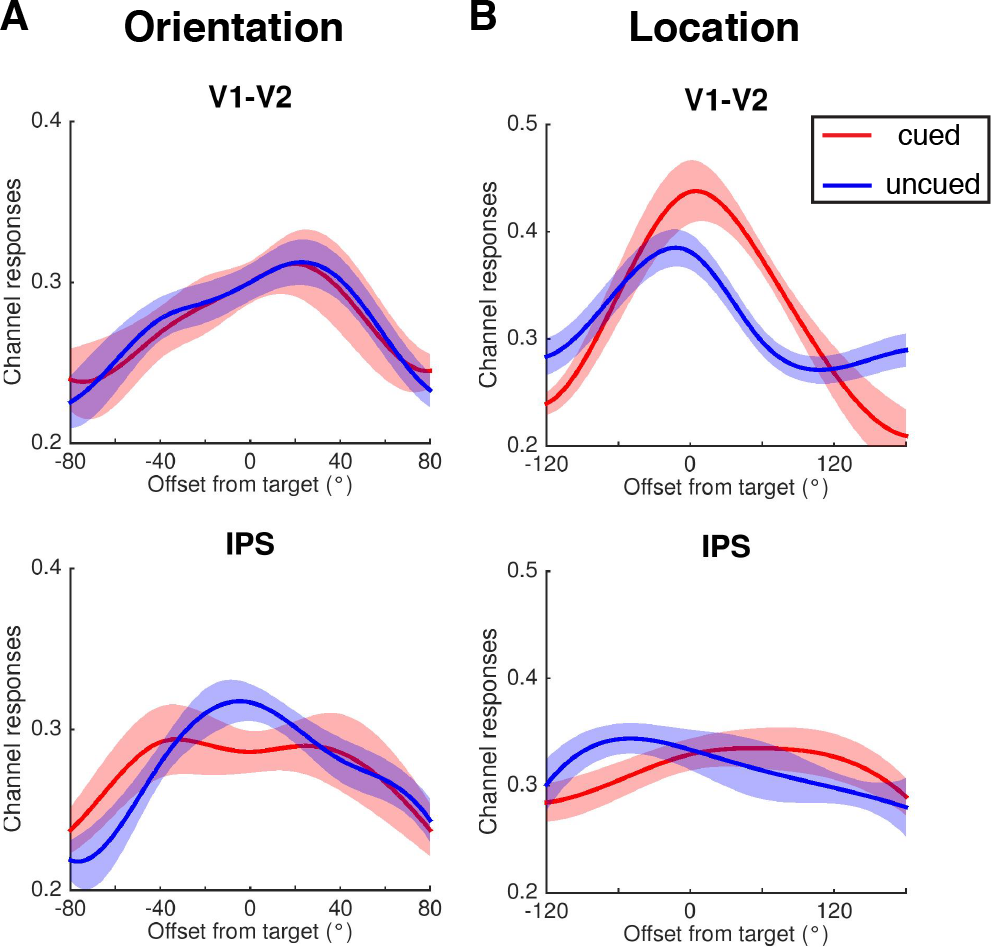
Experiment 2: Orientation and location reconstructions from a 1-item DR-trained IEM, in early visual ROI and in IPS. Orientation (**A**) and location (**B**) reconstructions during Delay1.2 in V1-V2 (early visual cortex) and IPS (parietal cortex), using a 1-item DR-trained model (trained from 4 s after trial onset for orientation and from 10 s after trial onset for location). Red line represents the cued orientation (AMI during Delay1.2), and blue line represents the uncued orientation (UMI during Delay1.2). Reconstructions were averaged across all participants. Continuous curves were created with spline interpolation method for demonstration purposes. Channel responses are estimated BOLD responses in relative amplitude. Shaded areas indicate ± 1 SEM.

**S4 Fig.**
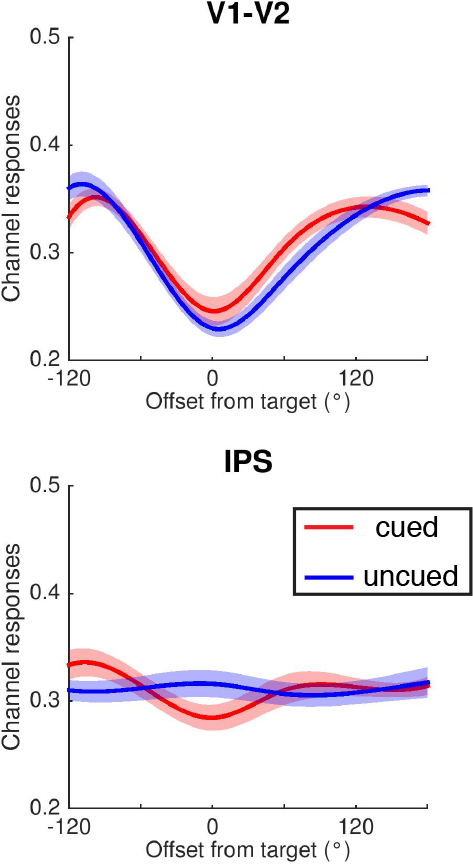
Experiment 2: Location reconstructions from a 1-item DR-trained IEM (sample-period data), in early visual ROI and in IPS. Location reconstructions during Delay1.2 in V1-V2 (early visual cortex) and IPS (parietal cortex), using a 1-item DR-trained model (trained from 4 s after trial onset*). Red line represents the cued orientation (AMI during Delay1.2), and blue line represents the uncued orientation (UMI during Delay1.2). Reconstructions were averaged across all participants. Continuous curves were created with spline interpolation method for demonstration purposes. Channel responses are estimated BOLD responses in relative amplitude. Shaded areas indicate ± 1 SEM. Although both the locations of the AMI and UMI could be reconstructed from sample-evoked data of 1-item DR task in early visual cortex, the pattern was negative reconstructions of both; S4 Fig); while AMI and UMI reconstructions from delay-period data of 1-item DR task were both positive (S3B Fig). Although not the focus of our current study, this result might suggest a similar mechanism [1] as what we reported in the main text: the brain recodes delay-period representations from those in the sample period so that information held in working memory can be protected from interference.

**S5 Fig.**
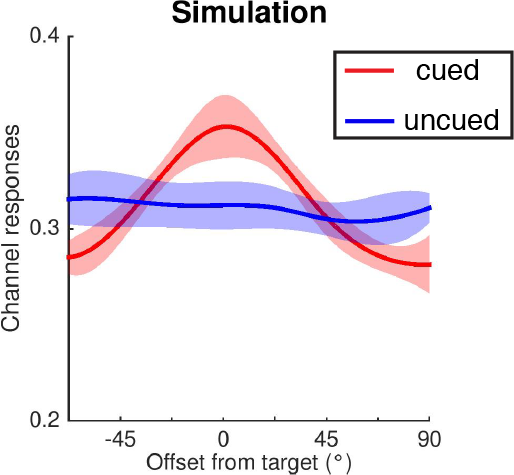
Reconstructions from a simulated IEM. The simulated IEM has eight orientation channels as in Experiment 1. Red line represents the simulated cued orientation (AMI during Delay1.2), and blue line represents the simulated uncued orientation (UMI during Delay1.2). Reconstructions were averaged across ten simulations. Continuous curves were created with spline interpolation method for demonstration purposes. Channel responses are estimated responses in relative amplitude. Shaded areas indicate ± 1 SEM.

**S1 Table.** *P*-values of amplitude from exponentiated cosine function fits

977 **Experiment 1:**
|  | **Retain1-cued** | **Retain1-uncued** | **Retain2-cued** | **Retain2-uncued** |
| --- | --- | --- | --- | --- |
| **V1-V2** | 0.038 | 0.180 | 0.168 | 0.038 |

**Experiment 2:**
|  | **Retain2-cued** | **Retain2-uncued** |
| --- | --- | --- |
| **Orientation:** |  |  |
| **V1-V2** | 0.010 | 0.049 |
| **IPS** | 0.029 | 0.075 |
| **Location:** |  |  |
| **V1-V2** | <0.00001 | 0.328 |
| **IPS** | 0.0004 | 0.035 |

All statistics were calculated for AMI-trained IEMs.

## References

1. Miller EK, Cohen JD. An integrative theory of prefrontal cortex function. Annu Rev Neurosci. 2001;24:167–202. Epub 2001/04/03. doi: 10.1146/annurev.neuro.24.1.167. PubMed PMID: 11283309.

2. Stokes MG, Kusunoki M, Sigala N, Nili H, Gaffan D, Duncan J. Dynamic coding for cognitive control in prefrontal cortex. Neuron. 2013;78(2):364–75. Epub 2013/04/09. doi: 10.1016/j.neuron.2013.01.039. PubMed PMID: 23562541; PubMed Central PMCID: PMCPMC3898895.

3. Baddeley A. Working memory: looking back and looking forward. Nat Rev Neurosci. 2003;4(10):829–39. Epub 2003/10/03. doi: 10.1038/nrn1201. PubMed PMID: 14523382.

4. D’Esposito M, Postle BR. The cognitive neuroscience of working memory. Annu Rev Psychol. 2015;66:115–42. Epub 2014/09/25. doi: 10.1146/annurev-psych-010814-015031. PubMed PMID: 25251486; PubMed Central PMCID: PMCPMC4374359.

5. Cowan N. Attention and memory: an integrated framework. New York: Oxford University Press; 1995. xv, 321 pages p.

6. Oberauer K. Access to information in working memory: exploring the focus of attention. J Exp Psychol Learn Mem Cogn. 2002;28(3):411–21. Epub 2002/05/23. PubMed PMID: 12018494.

7. Myers NE, Stokes MG, Nobre AC. Prioritizing Information during Working Memory: Beyond Sustained Internal Attention. Trends Cogn Sci. 2017;21(6):449–61. Epub 2017/04/30. doi: 10.1016/j.tics.2017.03.010. PubMed PMID: 28454719.

8. Sprague TC, Ester EF, Serences JT. Restoring Latent Visual Working Memory Representations in Human Cortex. Neuron. 2016;91(3):694–707. Epub 2016/08/09. doi: 10.1016/j.neuron.2016.07.006. PubMed PMID: 27497224; PubMed Central PMCID: PMCPMC4978188.

9. Owen AM, McMillan KM, Laird AR, Bullmore E. N-back working memory paradigm: a meta-analysis of normative functional neuroimaging studies. Hum Brain Mapp. 2005;25(1):46–59. Epub 2005/04/23. doi: 10.1002/hbm.20131. PubMed PMID: 15846822.

10. Conway AR, Kane MJ, Bunting MF, Hambrick DZ, Wilhelm O, Engle RW. Working memory span tasks: A methodological review and user’s guide. Psychon Bull Rev. 2005;12(5):769–86. Epub 2006/03/10. PubMed PMID: 16523997.

11. Larocque JJ, Lewis-Peacock JA, Postle BR. Multiple neural states of representation in short-term memory? It’s a matter of attention. Front Hum Neurosci. 2014;8:5. Epub 2014/01/31. doi: 10.3389/fnhum.2014.00005. PubMed PMID: 24478671; PubMed Central PMCID: PMCPMC3899521.

12. Lewis-Peacock JA, Drysdale AT, Oberauer K, Postle BR. Neural Evidence for a Distinction between Short-term Memory and the Focus of Attention. Journal of Cognitive Neuroscience. 2012;24(1):61–79. PubMed PMID: WOS:000297324600006.

13. LaRocque JJ, Lewis-Peacock JA, Drysdale AT, Oberauer K, Postle BR. Decoding attended information in short-term memory: an EEG study. J Cogn Neurosci. 2013;25(1):127–42. Epub 2012/12/04. doi: 10.1162/jocn_a_00305. PubMed PMID: 23198894; PubMed Central PMCID: PMCPMC3775605.

14. LaRocque JJ, Riggall AC, Emrich SM, Postle BR. Within-Category Decoding of Information in Different Attentional States in Short-Term Memory. Cereb Cortex. 2017;27(10):4881–90. Epub 2016/10/06. doi: 10.1093/cercor/bhw283. PubMed PMID: 27702811.

15. Rose NS, LaRocque JJ, Riggall AC, Gosseries O, Starrett MJ, Meyering EE, et al. Reactivation of latent working memories with transcranial magnetic stimulation. Science. 2016;354(6316):1136–9. Epub 2016/12/10. doi: 10.1126/science.aah7011. PubMed PMID: 27934762; PubMed Central PMCID: PMCPMC5221753.

16. Christophel TB, Iamshchinina P, Yan C, Allefeld C, Haynes JD. Cortical specialization for attended versus unattended working memory. Nat Neurosci. 2018;21(4):494–6. Epub 2018/03/07. doi: 10.1038/s41593-018-0094-4. PubMed PMID: 29507410.

17. Barak O, Tsodyks M. Working models of working memory. Curr Opin Neurobiol. 2014;25:20–4. Epub 2014/04/09. doi: 10.1016/j.conb.2013.10.008. PubMed PMID: 24709596.

18. Wolff MJ, Jochim J, Akyurek EG, Stokes MG. Dynamic hidden states underlying working-memory-guided behavior. Nat Neurosci. 2017;20(6):864–71. Epub 2017/04/18. doi: 10.1038/nn.4546. PubMed PMID: 28414333; PubMed Central PMCID: PMCPMC5446784.

19. Brouwer GJ, Heeger DJ. Cross-orientation suppression in human visual cortex. J Neurophysiol. 2011;106(5):2108–19. Epub 2011/07/22. doi: 10.1152/jn.00540.2011. PubMed PMID: 21775720; PubMed Central PMCID: PMCPMC3214101.

20. Brouwer GJ, Heeger DJ. Decoding and reconstructing color from responses in human visual cortex. J Neurosci. 2009;29(44):13992–4003. Epub 2009/11/06. doi: 10.1523/JNEUROSCI.3577-09.2009. PubMed PMID: 19890009; PubMed Central PMCID: PMCPMC2799419.

21. Sprague TC, Adam KCS, Foster JJ, Rahmati M, Sutterer DW, Vo VA. Inverted Encoding Models Assay Population-Level Stimulus Representations, Not Single-Unit Neural Tuning. eNeuro. 2018;5(3). Epub 2018/06/08. doi: 10.1523/ENEURO.0098-18.2018. PubMed PMID: 29876523; PubMed Central PMCID: PMCPMC5987635.

22. Ester EF, Sprague TC, Serences JT. Parietal and Frontal Cortex Encode Stimulus-Specific Mnemonic Representations during Visual Working Memory. Neuron. 2015;87(4):893–905. Epub 2015/08/11. doi: 10.1016/j.neuron.2015.07.013. PubMed PMID: 26257053; PubMed Central PMCID: PMCPMC4545683.

23. Xu Y. Reevaluating the Sensory Account of Visual Working Memory Storage. Trends Cogn Sci. 2017;21(10):794–815. Epub 2017/08/05. doi: 10.1016/j.tics.2017.06.013. PubMed PMID: 28774684.

24. Lorenc ES, Sreenivasan KK, Nee DE, Vandenbroucke ARE, D’Esposito M. Flexible coding of visual working memory representations during distraction. J Neurosci. 2018. Epub 2018/05/10. doi: 10.1523/JNEUROSCI.3061-17.2018. PubMed PMID: 29739867.

25. Oberauer K, Lin HY. An interference model of visual working memory. Psychol Rev. 2017;124(1):21–59. Epub 2016/11/22. doi: 10.1037/rev0000044. PubMed PMID: 27869455.

26. Schneegans S, Bays PM. Neural Architecture for Feature Binding in Visual Working Memory. J Neurosci. 2017;37(14):3913–25. Epub 2017/03/09. doi: 10.1523/JNEUROSCI.3493-16.2017. PubMed PMID: 28270569; PubMed Central PMCID: PMCPMC5394900.

27. Gosseries O, Yu Q, LaRocque JJ, Starrett MJ, Rose NS, Cowan N, et al. Parietal-Occipital Interactions Underlying Control- and Representation-Related Processes in Working Memory for Nonspatial Visual Features. J Neurosci. 2018;38(18):4357–66. Epub 2018/04/11. doi: 10.1523/JNEUROSCI.2747-17.2018. PubMed PMID: 29636395.

28. Mendoza-Halliday D, Torres S, Martinez-Trujillo JC. Sharp emergence of feature-selective sustained activity along the dorsal visual pathway. Nat Neurosci. 2014;17(9):1255–62. Epub 2014/08/12. doi: 10.1038/nn.3785. PubMed PMID: 25108910; PubMed Central PMCID: PMCPMC4978542.

29. Yu Q, Shim WM. Occipital, parietal, and frontal cortices selectively maintain task-relevant features of multi-feature objects in visual working memory. Neuroimage. 2017;157:97–107. Epub 2017/06/01. doi: 10.1016/j.neuroimage.2017.05.055. PubMed PMID: 28559190.

30. Serences JT, Ester EF, Vogel EK, Awh E. Stimulus-specific delay activity in human primary visual cortex. Psychol Sci. 2009;20(2):207–14. Epub 2009/01/28. doi: 10.1111/j.1467-9280.2009.02276.x. PubMed PMID: 19170936; PubMed Central PMCID: PMCPMC2875116.

31. Foster JJ, Bsales EM, Jaffe RJ, Awh E. Alpha-Band Activity Reveals Spontaneous Representations of Spatial Position in Visual Working Memory. Curr Biol. 2017;27(20):3216–23 e6. Epub 2017/10/17. doi: 10.1016/j.cub.2017.09.031. PubMed PMID: 29033335; PubMed Central PMCID: PMCPMC5661984.

32. Postle BR, Awh E, Serences JT, Sutterer DW, D’Esposito M. The positional-specificity effect reveals a passive-trace contribution to visual short-term memory. PLoS One. 2013;8(12):e83483. Epub 2014/01/05. doi: 10.1371/journal.pone.0083483. PubMed PMID: 24386212; PubMed Central PMCID: PMCPMC3873305.

33. Rajsic J, Wilson DE. Asymmetrical access to color and location in visual working memory. Atten Percept Psychophys. 2014;76(7):1902–13. Epub 2014/09/06. doi: 10.3758/s13414-014-0723-2. PubMed PMID: 25190322.

34. Sereno AB, Amador SC. Attention and memory-related responses of neurons in the lateral intraparietal area during spatial and shape-delayed match-to-sample tasks. J Neurophysiol. 2006;95(2):1078–98. Epub 2005/10/14. doi: 10.1152/jn.00431.2005. PubMed PMID: 16221750.

35. Bisley JW, Goldberg ME. Attention, intention, and priority in the parietal lobe. Annu Rev Neurosci. 2010;33:1–21. Epub 2010/03/03. doi: 10.1146/annurev-neuro-060909-152823. PubMed PMID: 20192813; PubMed Central PMCID: PMCPMC3683564.

36. Jerde TA, Merriam EP, Riggall AC, Hedges JH, Curtis CE. Prioritized maps of space in human frontoparietal cortex. J Neurosci. 2012;32(48):17382–90. Epub 2012/12/01. doi: 10.1523/JNEUROSCI.3810-12.2012. PubMed PMID: 23197729; PubMed Central PMCID: PMCPMC3544526.

37. Sprague TC, Serences JT. Attention modulates spatial priority maps in the human occipital, parietal and frontal cortices. Nat Neurosci. 2013;16(12):1879–87. Epub 2013/11/12. doi: 10.1038/nn.3574. PubMed PMID: 24212672; PubMed Central PMCID: PMCPMC3977704.

38. Bettencourt KC, Xu Y. Decoding the content of visual short-term memory under distraction in occipital and parietal areas. Nat Neurosci. 2016; 19(1):150–7. Epub 2015/11/26. doi: 10.1038/nn.4174. PubMed PMID: 26595654; PubMed Central PMCID: PMCPMC4696876.

39. Yu Q, Shim WM. Temporal-Order-Based Attentional Priority Modulates Mnemonic Representations in Parietal and Frontal Cortices. Cereb Cortex. 2018. Epub 2018/08/21. doi: 10.1093/cercor/bhy184. PubMed PMID: 30124789.

40. Koyluoglu OO, Pertzov Y, Manohar S, Husain M, Fiete IR. Fundamental bound on the persistence and capacity of short-term memory stored as graded persistent activity. Elife. 2017;6. Epub 2017/09/08. doi: 10.7554/eLife.22225. PubMed PMID: 28879851; PubMed Central PMCID: PMCPMC5779315.

41. Shannon CE. A mathematical theory of communication. The Bell System Technical Journal. 1948;27(3):379–423. doi: 10.1002/j.1538-7305.1948.tb01338.x.

42. Kaufman MT, Churchland MM, Ryu SI, Shenoy KV. Cortical activity in the null space: permitting preparation without movement. Nat Neurosci. 2014;17(3):440–8. Epub 2014/02/04. doi: 10.1038/nn.3643. PubMed PMID: 24487233; PubMed Central PMCID: PMCPMC3955357.

43. Spaak E, Watanabe K, Funahashi S, Stokes MG. Stable and Dynamic Coding for Working Memory in Primate Prefrontal Cortex. J Neurosci. 2017;37(27):6503–16. Epub 2017/06/01. doi: 10.1523/JNEUROSCI.3364-16.2017. PubMed PMID: 28559375; PubMed Central PMCID: PMCPMC5511881.

44. Barron HC, Vogels TP, Behrens TE, Ramaswami M. Inhibitory engrams in perception and memory. Proc Natl Acad Sci U S A. 2017;114(26):6666–74. Epub 2017/06/15. doi: 10.1073/pnas.1701812114. PubMed PMID: 28611219; PubMed Central PMCID: PMCPMC5495250.

45. Brainard DH. The Psychophysics Toolbox. Spat Vis. 1997;10(4):433–6. Epub 1997/01/01. PubMed PMID: 9176952.

46. Pelli DG. The VideoToolbox software for visual psychophysics: transforming numbers into movies. Spat Vis. 1997;10(4):437–42. Epub 1997/01/01. PubMed PMID: 9176953.

47. Bays PM, Catalao RF, Husain M. The precision of visual working memory is set by allocation of a shared resource. J Vis. 2009;9(10):7 1–11. Epub 2009/10/09. doi: 10.1167/9.10.7. PubMed PMID: 19810788; PubMed Central PMCID: PMCPMC3118422.

48. Cox RW. AFNI:: software for analysis and visualization of functional magnetic resonance neuroimages. Comput Biomed Res. 1996;29(3):162–73. Epub 1996/06/01. PubMed PMID: 8812068.

49. Wang L, Mruczek REB, Arcaro MJ, Kastner S. Probabilistic maps of visual topography in human cortex. Cerebral Cortex. 2015;25:3911–31.

50. Samaha J, Sprague TC, Postle BR. Decoding and Reconstructing the Focus of Spatial Attention from the Topography of Alpha-band Oscillations. J Cogn Neurosci. 2016;28(8):1090–7. Epub 2016/03/24. doi: 10.1162/jocn_a_00955. PubMed PMID: 27003790; PubMed Central PMCID: PMCPMC5074376

## References

1. Linke AC, Vicente-Grabovetsky A, Cusack R. Stimulus-specific suppression preserves information in auditory short-term memory. Proc Natl Acad Sci U S A. 2011;108(31):12961–6. Epub 2011/07/20. doi: 10.1073/pnas.1102118108. PubMed PMID: 21768383; PubMed Central PMCID: PMCPMC3150893.

